# Emergence of Non-Linear Mixed Selectivity in Prefrontal Cortex after Training

**DOI:** 10.1101/2020.08.02.233247

**Authors:** Wenhao Dang, Russell J. Jaffe, Xue-Lian Qi, Christos Constantinidis

**Affiliations:** Department of Neurobiology & Anatomy, Wake Forest School of Medicine, Winston-Salem, NC 27157, USA

**Author notes:** Address correspondence to: Christos Constantinidis, PhD., Department of Neurobiology and Anatomy, Wake Forest School of Medicine, Medical Center Blvd, Winston Salem, NC, 27157. These authors contributed equally to this work (Wenhao Dang, Russell J. Jaffe).

## Abstract

Neurons in the prefrontal cortex (PFC) are typically activated by different cognitive tasks, and also by different stimuli and abstract variables within these tasks. A single neuron’s selectivity for a given stimulus dimension often changes depending on its context, a phenomenon known as nonlinear mixed selectivity (NMS). It has previously been hypothesized that NMS emerges as a result of training to perform tasks in different contexts. We tested this hypothesis directly by examining the neuronal responses of different PFC areas before and after monkeys were trained to perform different working memory tasks involving visual stimulus locations and/or shapes. We found that training induces a modest increase in the proportion of PFC neurons with NMS exclusively for spatial working memory, but not shape working memory tasks, with area 9/46 undergoing the most significant increase in NMS cell proportion. We also found that increased working memory task complexity, in the form of simultaneously storing location and shape combinations, does not increase the degree of NMS for stimulus shape with other task variables. Lastly, in contrast to the previous studies, we did not find evidence that NMS is predictive of task performance. Our results thus provide critical insights on the representation of stimuli and task information in neuronal populations, which may pave the way to a greater understanding of neural selectivity in working memory.

**SIGNIFICANCE STATEMENT:** How multiple types of information are represented in working memory remains a complex computational problem. It has been hypothesized that nonlinear mixed selectivity allows neurons to efficiently encode multiple stimuli in different contexts, after subjects have been trained in complex tasks. Our analysis of prefrontal recordings obtained before and after training monkeys to perform working memory tasks only partially agreed with this prediction, in that nonlinear mixed selectivity emerged for spatial but not shape information, and mostly in mid-dorsal PFC. Nonlinear mixed selectivity also displayed little modulation across either task complexity or correct performance. These results point to other mechanisms, in addition to nonlinear mixed selectivity, to represent complex information about stimulus and context in neuronal activity.

## INTRODUCTION

Working memory (WM) is broadly defined as the ability to encode, maintain, and manipulate information in the conscious mind over a period of seconds without the presence of any sensory inputs. As a core component of complex cognitive abilities such as planning and reasoning, the true importance of WM ultimately depends on whether it can maintain and manipulate task relevant information in a task relevant manner (Baddeley, 2012). Multiple variables, including external sensory inputs and internal task requirements, must be encoded in order to achieve the level of adaptability in WM that is necessary for complex tasks. The mechanisms that underlie this encoding process across time and neuronal population is one of the most important questions in current WM research.

When individuals are required to maintain objects in their WM, neurons from a network of brain regions may exhibit selective and sustained increases or decreases in their activity in order to represent the remembered objects through these unique patterns of activity (Constantinidis and Procyk, 2004). The prefrontal cortex (PFC) plays a leading role in this network, and by extension, in the use of WM (Riley and Constantinidis, 2016). For example, when the PFC is damaged or degraded, whether through trauma, illness, or experimental lesions, performance in WM tasks seems to decrease dramatically (Curtis and D’Esposito, 2004; Morris and Baddeley, 1988; Rossi et al., 2007).

Individual PFC neurons typically encode more than one variables, and the exact variables encoded are task dependent (Asaad et al., 2000; Machens et al., 2010; Mansouri et al., 2006; Qi et al., 2015; Warden and Miller, 2010). More interestingly, a portion of neurons exhibit nonlinear mixed selectivity (NMS) for different variables, which means that their response to the combination of variables cannot be predicted by the linear summation of their responses to single variables (Johnston et al., 2020; Parthasarathy et al., 2017; Rigotti et al., 2013). Theoretical studies have shown that NMS is useful for linear readouts of flexible, arbitrary combinations of variables (Buonomano and Maass, 2009; Fusi et al., 2016; Rigotti et al., 2010), and may also control the trade-off between discrimination and generalization (Barak et al., 2013; Johnston et al., 2020).

Despite the proposed importance of NMS on theoretical grounds, some experimental studies have failed to detect neurons with NMS (Cavanagh et al., 2018). It is therefore possible that NMS may manifest exclusively in a limited set of PFC subdivisions or alternatively, that NMS emerges exclusively after training to perform specific types of cognitive tasks. Moreover, the implications of NMS on other aspects of neural encoding, such as code stability, have not yet been investigated. We were therefore motivated to analyze and compare neural data from rhesus macaque monkeys before and after training. Here we report results of NMS as a function of task training, performance of different types of working memory tasks, and correct and error trials, across different prefrontal areas.

## METHODS

### Animals

Data obtained from six male rhesus monkeys (*Macaca mulatta*), age 5–9 years old, weighing 5–12 kg, as previously documented (Riley et al., 2018), were analyzed in this study. None of the animals had any prior experimentation experience at the onset of our study. Monkeys were either single-housed or pair-housed in communal rooms with sensory interactions with other monkeys. All experimental procedures followed guidelines set by the U.S. Public Health Service Policy on Humane Care and Use of Laboratory Animals and the National Research Council’s Guide for the Care and Use of Laboratory Animals and were reviewed and approved by the Wake Forest University Institutional Animal Care and Use Committee.

### Experimental setup

Monkeys sat with their head fixed in a primate chair while viewing a monitor positioned 68 cm away from their eyes with dim ambient illumination. Animals were required to fixate on a 0.2° white square appearing in the center of the screen. During each trial, the animals maintained fixation on the square while visual stimuli were presented either at a peripheral location or over the fovea in order to receive a juice reward. Any break of fixation immediately terminated the trial and no reward was given. Eye position was monitored throughout the trial using a non-invasive, infrared eye position scanning system (model RK-716; ISCAN, Burlington, MA). The system achieved a < 0.3° resolution around the center of vision. Eye position was sampled at 240 Hz, digitized and recorded. Visual stimuli display, monitoring of eye position, and the synchronization of stimuli with neurophysiological data were performed with in-house software implemented on the MATLAB environment (Mathworks, Natick, MA), and utilizing the Psychophysics Toolbox (Meyer and Constantinidis, 2005).

### Pre-training task

Following a brief period of fixation training and acclimation to the stimuli, monkeys were required to fixate on a center position while stimuli were displayed on the screen. The monkeys were rewarded for maintaining fixation during the trial with a liquid reward (fruit juice). The stimuli shown in the pre-training passive spatial task were white 2° squares, presented in one of nine possible locations arranged in a 3 × 3 grid with 10° distance between adjacent stimuli. The stimuli shown in the pre-training passive feature task were white 2° circles, diamonds, H-letters, hashtags, plus signs, squares, triangles, or inverted Y-letters, also presented in one of nine possible locations arranged in a 3 × 3 grid with 10° distance between adjacent stimuli.

Presentation began with a fixation interval of 1 s where only the fixation point was displayed, followed by a 500 ms of stimulus presentation (referred to hereafter as cue), followed by a 1.5 s “delay” interval (referred to hereafter as delay1) where, again, only the fixation point was displayed. A second stimulus (referred to hereafter as sample) was subsequently shown for 500 ms. In the spatial task, this second stimulus would be either identical in location to the initial stimulus, or diametrically opposite the first stimulus. In the feature task, this second stimulus would always be identical in location to the initial stimulus and would either be an identical shape or the corresponding non-match shape (each shape was paired with one non-match shape).

In both the spatial and feature task, this second stimulus display was followed by another “delay” period (referred to hereafter as delay2) of 1.5 s where only the fixation point was displayed. The location and identity of stimuli was of no behavioral relevance to the monkeys during the “pre-training” phase, as fixation was the only necessary action for obtaining reward.

### Post-training task

Four of the six monkeys were trained to complete active spatial, feature and conjunction WM tasks. These active spatial and feature tasks were identical to the passive spatial and feature tasks that were applied during the “pre-training” phase, except that these tasks now required the monkeys to remember the spatial location or the shape feature of the first presented stimulus, and report whether the second stimulus matched the spatial location or shape feature of the first stimulus, respectively, via saccading to one of two target stimuli (green for match, blue for non-match). Each target stimulus could appear at one of two locations orthogonal to the cue/sample stimuli, pseudo-randomized in each trial.

The conjunction task combined the active spatial and feature tasks, and the stimuli shown were a white 2° circle, diamond, H-letter, hashtag, plus sign, square, triangle, or inverted Y-letter shapes, presented in one of nine possible locations arranged in a 3 × 3 grid with 10° distance between adjacent stimuli. Each trial consisted of a fixation interval of 1 s where only the fixation point was displayed, followed by 500 ms of the first stimulus presentation, followed by a 1.5 s delay interval where, again, only the fixation point was displayed. A second stimulus was subsequently shown for 500 ms, and after a second, 1.5 s delay period the monkeys would report whether the second stimulus matched both the spatial location and shape feature of the first stimulus, via saccading to one of the two target stimuli. The conjunction task was therefore the most complex task, as the monkeys were required to simultaneously store two different items—location and shape—in their WM.

### Surgery and neurophysiology

A 20 mm diameter craniotomy was performed over the PFC and a recording cylinder was implanted over the site. The location of the cylinder was visualized through anatomical magnetic resonance imaging (MRI) and stereotaxic coordinates post-surgery. For two of the four monkeys the recording cylinder was moved after an initial round of recordings in the post-training phase to sample an additional surface of the PFC.

### Anatomical localization

Each monkey underwent an MRI scan prior to neurophysiological recordings. Electrode penetrations were mapped onto the cortical surface. We identified 6 lateral PFC regions: a posterior-dorsal region that included area 8A, a mid-dorsal region that included area 8B and area 9/ 46, an anterior-dorsal region that included area 9 and area 46, a posterior-ventral region that included area 45, an anterior-ventral region that included area 47/12, and a frontopolar region that included area 10. However, the frontopolar region was not sampled sufficiently to be included in the present analyses.

In addition to comparisons between brain areas segmented in this fashion, other analyses were performed to account for the position of each neuron along the AP axis. For the purposes of our analysis, we defined the AP axis as the line connecting the genu of the arcuate sulcus to the frontal pole. The recording coordinates of each neuron were projected onto this line, with position expressed as a proportion of the line’s length.

### Neuronal recordings

Neural recordings were carried out in areas 8, 9, 9/46, 45, 46, and 47/12 of the PFC both before and after training in each WM task. Subsets of the data presented here were previously used to determine the collective properties of neurons in the dorsal and ventral PFC, as well as the properties of neurons before and after training in the posterior-dorsal, mid-dorsal, anterior-dorsal, posterior-ventral, and anterior-ventral PFC subdivisions. Extracellular recordings were performed with multiple microelectrodes that were either glass- or epoxylite-coated tungsten, with a 250 μm diameter and 1–4 MΩ impedance at 1 kHz (Alpha-Omega Engineering, Nazareth, Israel). A Microdrive system (EPS drive, Alpha-Omega Engineering) advanced arrays of up to 8-microelectrodes, spaced 0.2–1.5 mm apart, through the dura and into the PFC. The signal from each electrode was amplified and band-pass filtered between 500 Hz and 8 kHz while being recorded with a modular data acquisition system (APM system, FHC, Bowdoin, ME). Waveforms that exceeded a user-defined threshold were sampled at 25 μs resolution, digitized, and stored for off-line analysis. Neurons were sampled in an unbiased fashion, collecting data from all units isolated from our electrodes, with no regard to the response properties of the isolated neurons. A semi-automated cluster analysis relied on the KlustaKwik algorithm, which applied principal component analysis of the waveforms, to sort recorded spike waveforms into separate units. To ensure a stable firing rate in the analyzed recordings, we identified recordings in which a significant effect of trial sequence was evident at the baseline firing rate (ANOVA, p < 0.05), e.g., due to a neuron disappearing or appearing during a run, as we were collecting data from multiple electrodes. Data from these sessions were truncated so that analysis was only performed on a range of trials with stable firing rate. Less than 10% of neurons were corrected in this way. Identical data collection procedures, recording equipment, and spike sorting algorithms were used before and after training in order to prevent any analytical confounds.

### Data analysis

Data analysis was implemented with the MATLAB computational environment (Mathworks, Natick, MA), with additional statistic tests implemented through Originlab (OriginLab Corporation, Northampton, MA) and StatsDirect (StatsDirect Ltd. England). Peristimulus time histograms (PSTHs) for illustrations were calculated through the moving window average method with a Gaussian window that had a 200 ms standard deviation, with the shaded area indicating two times standard error cross trials. For all tasks, only cells with at least 12 correct trials for each cue-sample location/shape pairs were included in the analysis. To classify neurons of the spatial task into different categories of selectivity, we performed two-way ANOVAs on the spike count between either the stimuli location x matching status in for the trial, or between stimuli location x task epoch (first or second stimulus presentation). Neurons with classic selectivity (CS) exhibited a main effect of only one factor without significant interaction term. Neurons with linear mixed selectivity (LMS) exhibited main effects of both factors without significant interaction term. Neurons with NMS exhibited a significant interactions term. Finally, non-selective (NS) neurons exhibited no significant term for both the main effects and the interaction term. Similarly, the two factors for feature task ANOVA analysis were stimuli shape x matching status, and stimuli shape x task epoch for the trial.

A method based on singular value decomposition (SVD) and cross validation was used to calculate population dimensionality for the spatial and feature tasks (Ahlheim and Love, 2018). The dimensionality of a matrix is defined as its number of non-zero singular values, identified by SVD. Under this condition however, the noise in the recorded neural data could potentially inflate the number of non-zero singular values, even if the true dimensionality were low. Reconstruction from components with cross-validation could be used to estimate dimensionality with noise, under the assumption that only true underlying dimensionality can contribute to the reconstruction performance in the cross-validation dataset. In short, data from j trials were randomly assigned to training the dataset with j-2 trials, using one of the remaining trials for validation and the other for testing. SVD was then applied to the averaged training data to obtain all of the possible low-dimensional reconstructions, which were then correlated through a validation run. The dimensionality that produced the highest average correlation across j-1 runs was selected as dimensionality estimate k for this fold, and a k-dimensional reconstruction was correlated with the held-out test data, resulting in the final reconstruction correlation. A similar method had also been recently used to estimate the dimensionality over time for neural data (Cueva et al., 2020). The dimensionality of the sample and delay2 period in the spatial and feature task was calculated on 50 resamples of a 200-cell pseudo-population in the corresponding datasets.

Only PFC areas with more than 50 cells in both pre- and post-training time points were included in the subdivision mixed selectivity comparison analysis. Thus, for the feature task, only data from the mid-dorsal, posterior-dorsal and posterior-ventral PFC were analyzed, while the spatial task analyzed data from the mid-dorsal, posterior-dorsal, posterior-ventral, anterior-dorsal and anterior-ventral PFC.

Neural data from tasks that applied the exact same visual stimuli were used to compare mixed selectivity between feature/spatial and conjunction tasks. For example, to compare the feature and the conjunction tasks, we started by selecting a subset of conjunction trials, in which both visual stimuli appeared at the same location as the corresponding feature task trials. Then, since all eight shapes were used in a single recording session for the feature task, a subset of trials, in which same shape pairs were used as the corresponding conjunction task, could be chosen as the feature dataset. Our prior methods of ANOVA analyses could thus be applied for comparison across these datasets.

For comparing mixed selectivity in success and error trials from the spatial task, we first examined two task variables—the stimuli location and matching status. In this analysis, we utilized neurons that had at least 3 match and non-match trials for both the correct and error dataset, in at least 3 stimuli locations. The number of minimum trials and stimuli locations were chosen to maximize the average trial number for each cell included into the analysis, while still retaining a sufficiently large sample (i.e. >150 cells). The same number of trials from each stimuli location were randomly chosen in the correct and error dataset. This randomized trial selection process was repeated 50 times in order to make the best use of the uneven number of available trials in two datasets. We also analyzed factors of stimuli location and task epoch. For this analysis, we used neural data from match trials only, and from neurons that had at least 4 correct and error trials, in at least 4 stimuli locations.

For decoding analysis, spiking responses from 1 second before cue onset to 5 seconds after cue onset were first binned using a 400 ms wide window and 100 ms steps to create a spike count vector with a length of 57 elements. A pseudo-population was then constructed using the spike count vectors from all the available neurons of all the available animals, thus resulting in a dataset with 96 trials, as if they were recorded simultaneously. The population response matrix was z-score normalized before being used to train the decoder. A linear Support Vector Machine (SVM) decoding algorithm was implemented using the MATLAB fitcecoc function to decode stimuli location, stimuli shape, or the match/non-match status of trials. A 10-fold cross validation method was used to estimate the decoder performance and 20 random samplings were implemented to calculate a 95% confidence interval. For the location and feature task, the decoding baseline for sensory information was 12.5%, since there were 8 different options, and 50% for the matching status, since there were only 2 different options. In the pre-training vs post-training decoding analysis (Fig. 8), linear (CS and LMS) and nonlinear (NMS) neurons are first defined by their pre and post training responses in the sample or delay2 period. Each classified population was then applied to decode sensory information (location and shape) and matching status. A randomization test was used to determine the time points at which decoding performance was significantly different between different selectivity categories. In short, we constructed the null distribution by randomly reassigning the cell selectivity labels under comparison, and re-computing the maximum absolute difference across all time points of the data in each iteration. This procedure was repeated for 5000 times. A difference was deemed to be significant if the true response difference occurred at the extremes of this null distribution (p <0.05, two-tailed). Since every point in the null distribution is the maximum of all time points, this method already corrected for the multiple comparisons.

Only informative neurons (CS, LMS, and NMS neurons) in the delay2 period were used for the cross temporal decoding analysis (Fig. 7), since we wanted to explore the decoding dynamics during the delay period. The linear SVM decoder was trained on individual time points and thus had 57 linear decision boundaries. The same dataset was then classified by every decision boundary in the vector to produce a 57×57 matrix—a process that was repeated 20 times in order to eliminate the noise. The decoding performance matrix for each condition was normalized individually to highlight the coding dynamics rather than absolute performance.

### Data availability

All relevant data and code will be available from the corresponding author on reasonable request. Matlab decoder code for figure 8 and 9 is available at https://github.com/dwhzlh87/mixed-selectivity

## RESULTS

Extracellular neurophysiological recordings were collected from the lateral PFC of six monkeys before and after they were trained to perform the match/nonmatch WM tasks (Meyer et al., 2011; Riley et al., 2018). The task required them to view two stimuli appearing in sequence, with delay periods intervening between them, and to report whether or not the second stimulus was identical to the first. The two stimuli could differ in terms of their location (spatial task, Fig. 1A), shape (feature task, Fig. 1B), or both (conjunction task, Fig. 1C). If the second stimulus matched with the first, monkeys would saccade towards a green target during a subsequent interval, otherwise to a blue target at a diametrical location. A total of 1617 cells from six monkeys and 1495 cells from five monkeys were recorded while the animals were performing the passive spatial and feature tasks, respectively, which were mutually dubbed “pre-training.” A total of 1104 cells from three monkeys and 1116 cells from two monkeys were collected while the animals were performing the active spatial and feature tasks, respectively, which were mutually dubbed “post-training”. We also collected neural data from 247 neurons for the passive spatial task from two monkeys after they were trained in the active spatial task.

**Figure 1.**
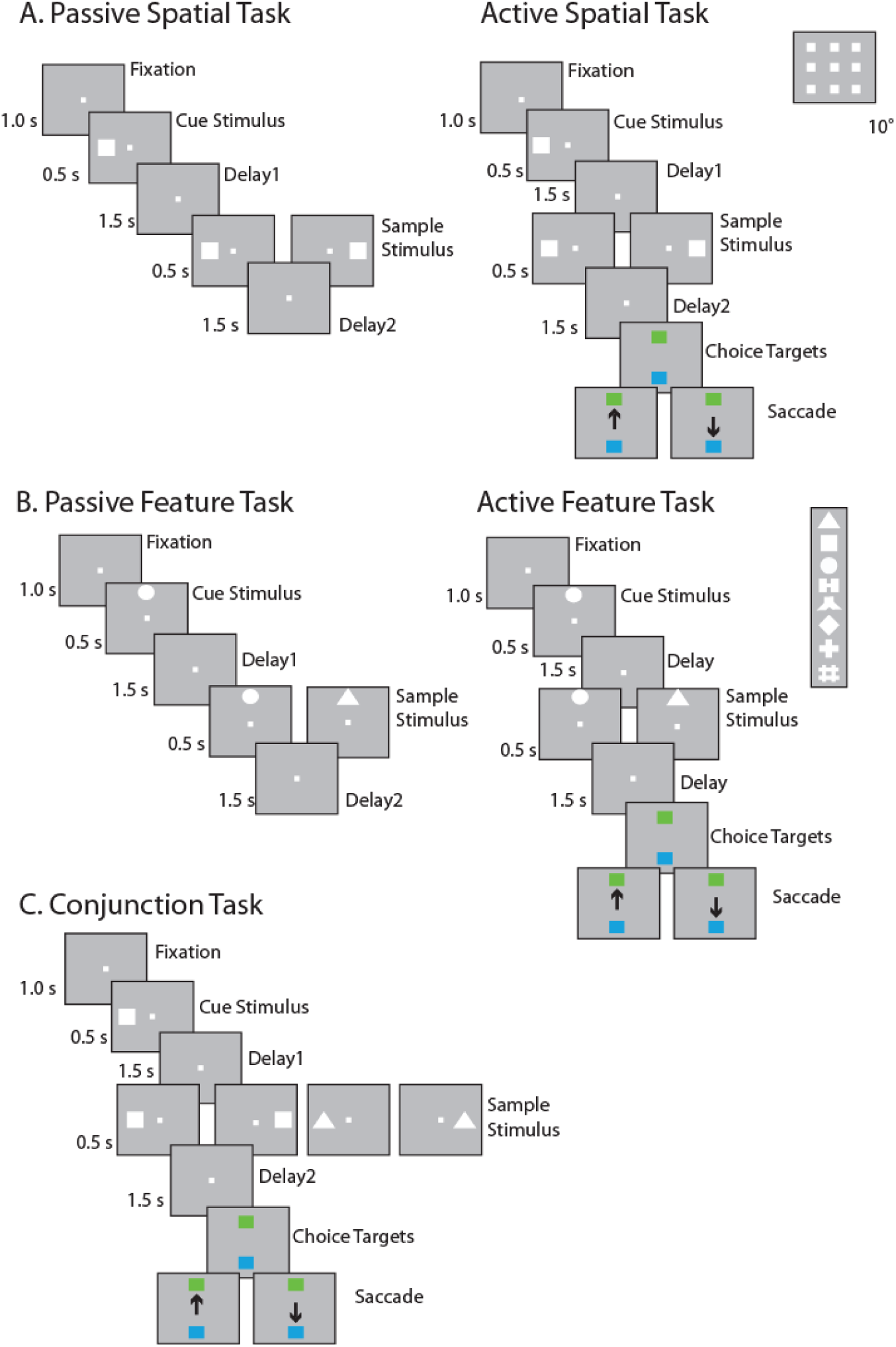
Task structure and stimuli used. The animals were required to maintain center fixation throughout both active and passive task trials. At the end of active tasks trials however, monkeys were required to make a saccade to a green target if the stimuli matched or to a blue target if the stimuli did not match. (A) Spatial location match-to-sample task, nine possible cue locations in a session shown in the inset. (B) Shape feature match-to-sample task, 8 possible shapes in a session shown in the inset. (C) Spatial-shape conjunction task, up to two locations and two stimuli shapes were used for any single particular session. Stimuli in all tasks extended 2 degree of visual angle.

An additional 975 cells from two monkeys were collected while they were performing the active “post-training” conjunction task.

### Types of selectivity in individual neuronal responses

In our tasks, the context of a given stimulus depends upon the task interval and sequence in which it is presented. We first considered how selectivity for stimulus location and shape in the spatial, feature and conjunction WM tasks may vary when the same sample stimulus appears as a match (it is preceded by a cue at the same location/shape) or a nonmatch (i.e. is preceded by a cue stimulus of a different location/shape). The neuronal firing rate is therefore a function of the stimulus location/shape (eight shapes, eight locations arranged on a 3×3 grid with 10 degrees distance between stimuli, excluding the center location) and whether this sample stimulus matched the cue stimulus. We used a 2-way ANOVA with the factors of stimulus location/shape and match/nonmatch status to classify neurons into four categories of selectivity. CS neurons exhibited a significant main effect to only one of the factors (stimulus identity or matching status) and had no significant interaction term. In Fig. 2, the first exemplar displays a cell selective exclusively for the location of the stimuli, which does not respond differently regardless of whether the stimulus appeared as a match or nonmatch. The second exemplar of Fig. 2 displays a cell not selective for the location of the stimuli but demonstrates higher mean response when the stimulus appeared as a non-match. LMS neurons exhibited a significant main effect for both factors but had no significant interaction term. The third exemplar of Fig. 2 displays a neuron demonstrating a higher mean response when stimuli appear as non-match, while simultaneously displaying the same rank order preference for location. NMS neurons exhibited a significant interaction effect, as shown in the last exemplar in Fig. 2, a neuron demonstrating different selectivity pattern for locations under match vs. non-match conditions. Finally, NS indicated the neurons with no selectivity to any factors or their interaction under consideration. These analyses were performed using the firing rate recorded during the stimulus presentation period, and again, using the firing rate recorded during the delay period that followed it.

**Figure 2.**
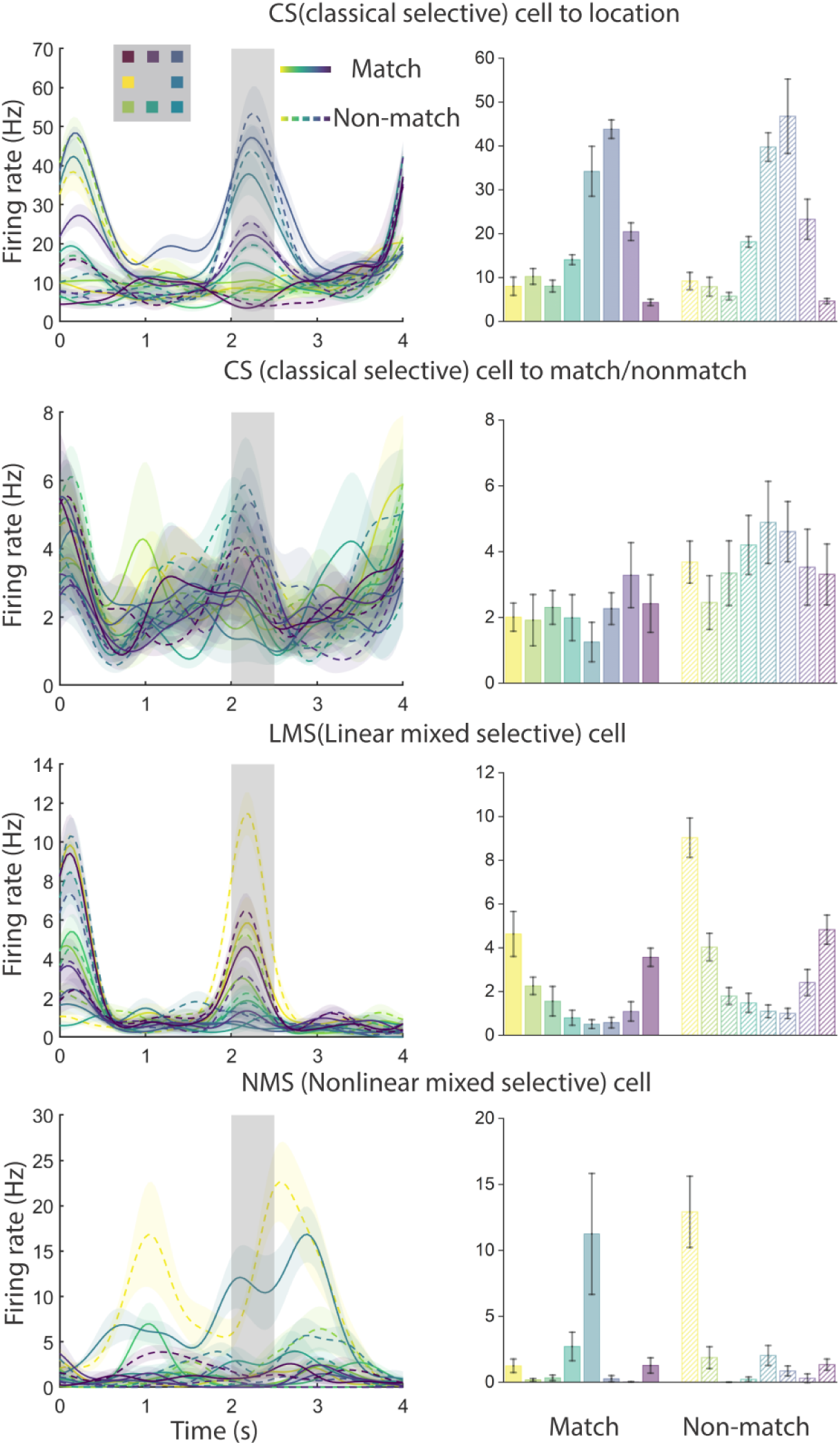
Exemplar neural responses from the spatial task for CS (classical selective), LMS (linear mixed selective) and NMS (nonlinear mixed selective) cells, defined by the task variables of stimulus location and match status. Selectivity classification were based on the spike responses of the 500 ms sample period. The locations the stimuli were color coded, with a solid line/bar representing when the stimulus was a match with the cue, and a dash line/bar representing when the stimulus was a nonmatch with the cue. Shaded regions and error bar indicate 2 times SE of firing rate.

A second type of NMS was identified in terms of selectivity for stimulus sequence, that is, whether the same stimulus appeared first (cue) or second (sample). To avoid the confound of the match or nonmatch status of the second stimulus, we relied exclusively on match stimuli. This form of NMS was also evaluated through a 2-way ANOVA model, identifying CS, LMS, NMS, and NS neurons in terms of how the neurons represented the exact same stimulus when it appeared as a cue and as a match stimulus.

### Effects of training on NMS

When we used the factors of stimulus location/shape and match/nonmatch status for our two-way ANOVA, we found that training in the spatial WM task increased the proportion of NMS cells in both the sample period and the delay period that followed the sample (sample period: pre-training proportion=6.2%, post-training proportion=12.3%, two-sample proportion test, z=5.31, p=1.13×10^−7^; delay2 period: pre-training proportion= 2.8%, post-training proportion=6.2%, two-sample proportion test, z=4.62, p=4.86×10^−5^). However, this increase in selectivity was not exclusive to NMS cells. The proportion of CS cells also increased in the delay period following the sample (pre-training proportion= 10.6%, post-training proportion=14.8%, two-sample proportion test, z=3.19, p=0.0014).

The increase in NMS cells was not evident for all types of training. When we looked at the proportion of change across the pre-training and post-training feature task, we only found an increase of proportion for CS cells (sample period: pre-training proportion= 12.0%, post-training proportion=15.7%, two-sample proportion test, z=2.65, p=0.0081; delay2 period: pre-training proportion= 9.0%, post-training proportion=22.6%, two-sample proportion test, z=9.37, p=0). No significant increase in the proportion of NMS cells was observed (sample period: pre-training proportion=5.8%, post-training proportion=6.7%, two-sample proportion test, z=1.01, p=0.314; delay2 period: pre-training proportion= 4.2%, post-training proportion=4.6%, two-sample proportion test, z=0.522, p=0.602) (Fig. 3 A,B).

**Figure 3.**
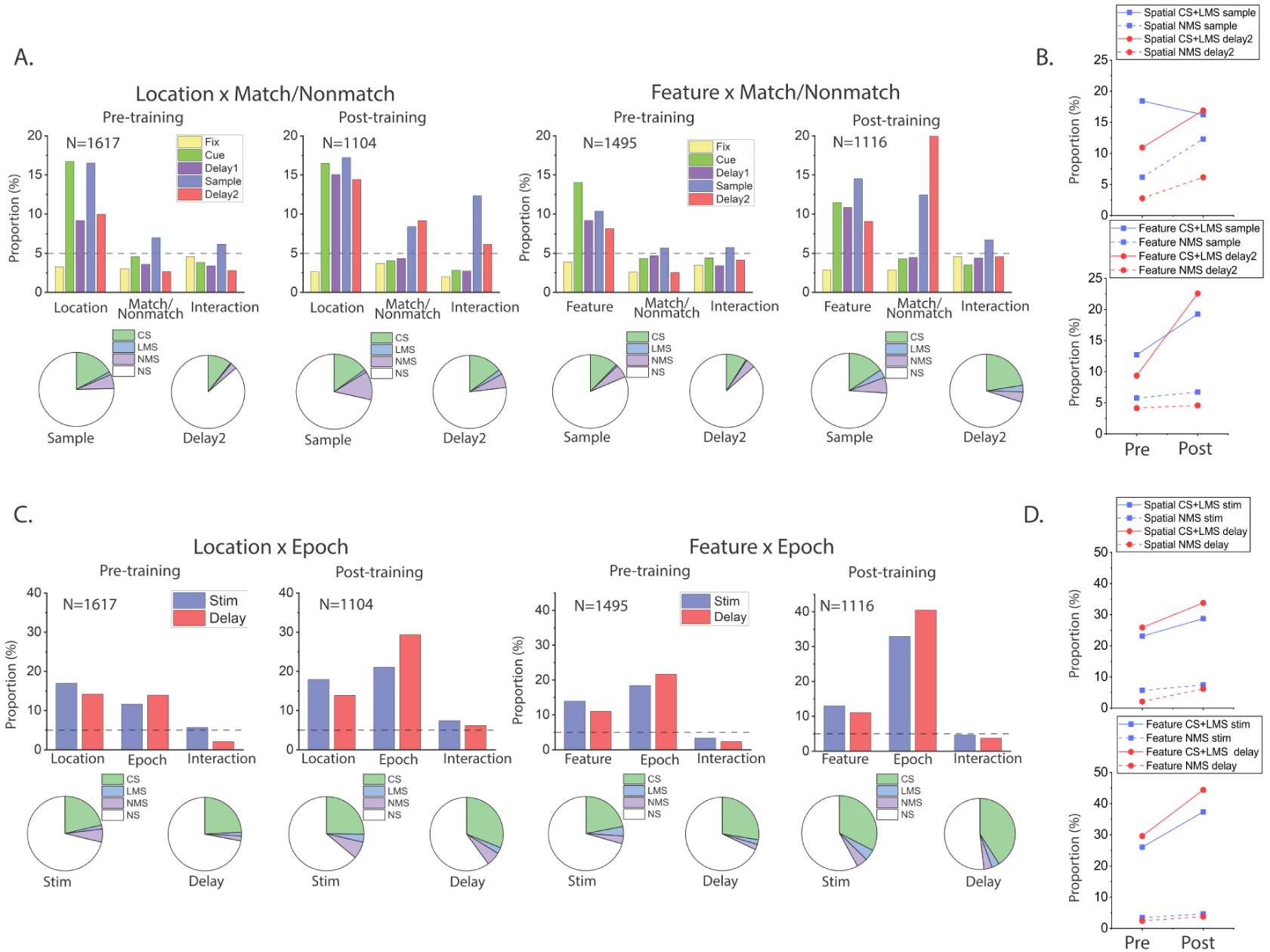
Training increased mixed selectivity preferentially in the spatial task. (A) Bar graphs show the proportions of cells tuned to stimuli identities (Location/Shape), matching status and their interaction (i.e. NMS) in different stages of the task trials, both before and after the animals were trained for the active tasks. Pie charts show the proportion of different selectivity categories (NS, CS, LMS and NMS) in the sample and delay2 periods of both tasks, both before and after the animals were trained for the active tasks. (B) Plots of corresponding proportion changes. (C) Same as (A) but examining the interaction between stimuli identities (Location/Shape), and task epoch (cue/delay1 vs sample/delay2 period), instead of trials matching status. (D) Plots of corresponding proportion changes.

Similar results were observed when we used the factors of stimulus location/shape and task epoch (cue vs. match) for the two-way ANOVA instead (Fig. 3 C,D). For the spatial task, training increased the proportion of NMS cells, at least in the delay period (stimulus period: pre-training proportion= 5.7%, post-training proportion=7.4%, two-sample proportion test, z=1.78, p=0.075; delay period: pre-training proportion= 2.1%, post-training proportion=6.2%, two-sample proportion test, z=5.03, p=4.94×10^−7^). A similar increase was observed for the CS cells (stimulus period: pre-training proportion= 21.3%, post-training proportion=25.3%, two-sample proportion test, z=2.37, p=0.018; delay period: pre-training proportion= 24.2%, post-training proportion=31.1%, two-sample proportion test, z=3.93, p=8.53×10^−5^). In the feature task, once again, only the proportion of CS cells changed (stimulus period: pre-training proportion= 21.9%, post-training proportion=32.6%, two-sample proportion test, z=6.05, p=1.45×10^−9^; delay period: pre-training proportion= 27.6%, post-training proportion=41%, two-sample proportion test, z=7.16, p=8.32×10^−13^). The proportion of NMS cells with an effect in the stimulus period remained relatively unchanged for the cue/match period (pre-training proportion= 3.4%, post-training proportion=4.7%, two-sample proportion test, z=1.59, p=0.112), as well as the delay period (pre-training proportion= 2.4%, post-training proportion=3.8%, two-sample proportion test, z=1.95, p=0.051).

To further validate our proportional measure for NMS and compare our results to previous research on NMS in the PFC, we plotted the F scores for the interaction term (i.e. stimulus identity × matching status) in both the spatial and the feature task (Fig. 4 A). We found that this measure of NMS for individual cells increased specifically for the spatial task, indicated by much higher F score values after training for the spatial task. We also measured the dimensionality of population responses in the sample and delay2 period for the spatial and feature task. Again, this analysis confirmed the results of our cell proportion measure (Fig. 4 B). For the spatial task, there was a significant increase of dimensionality after training (sample period: pre-training dimensionality= 5.72, post-training dimensionality=10.33, two-sample t test, t(98)=12.21, p=2.18×10^−21^; sample period: pre-training dimensionality= 3.25, post-training dimensionality=6.29, two-sample t test, t(98)=9.39, p=2.51×10^−15^). For the feature task however, no significance increase was observed in mean F score (Fig. 4A) or dimensionality (Fig 4B; sample period: pre-training dimensionality= 2.72, post-training dimensionality=2.73, two-sample t test, t(98)=0.027, p=0.978; sample period: pre-training dimensionality= 2.43, post-training dimensionality=2.11, two-sample t test, t(98)=3.49, p=7.29×10^−4^).

**Figure 4.**
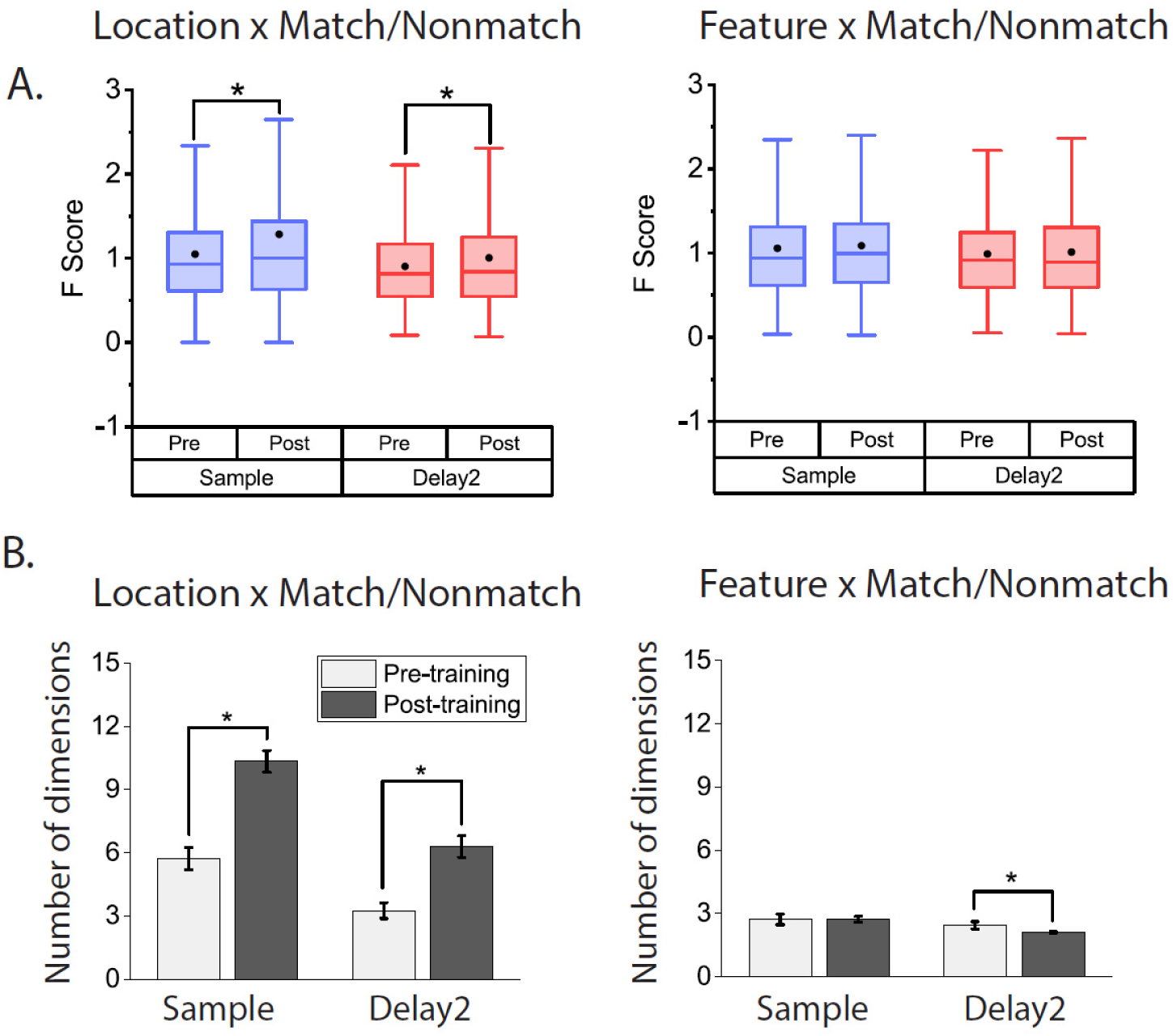
(A) Analysis of F score for the interaction term (stimuli identity × match/nonmatch) shows that the degree of mixed selectivity increased after training for the spatial task only. Black dots in the box represent mean, box boundaries indicate 25%-75% range, and whiskers represents 1.5 IQR. (B) Dimensionality measure of neural responses in the spatial (left) and feature (right) task, before and after training in the active tasks.

### Regional localization of NMS

To assess whether specific sub-regions of the PFC may be specialized for NMS, we divided the lateral PFC into six regions (Fig. 5 A) and analyzed the respective neurophysiological data from five of these regions in order to determine the different areas’ proportional contributions to the observed changes in NMS. We examined NMS defined by location/shape and match/nonmatch status in the sample period and ultimately found that the mid-dorsal subdivision underwent the greatest proportional change in NMS cells for the spatial task after training (Fig. 5 C), without a comparable increase in the proportion of CS neurons (Mid-dorsal: CS 21.7% pre-training to 19.0% post-training, NMS 8.2% pre-training to 21.8% post-training). For the feature task however, the proportional change in NMS cells was relatively small, with moderate increases in CS and LMS observed in all three analyzed areas (Fig. 5 B).

**Figure 5.**
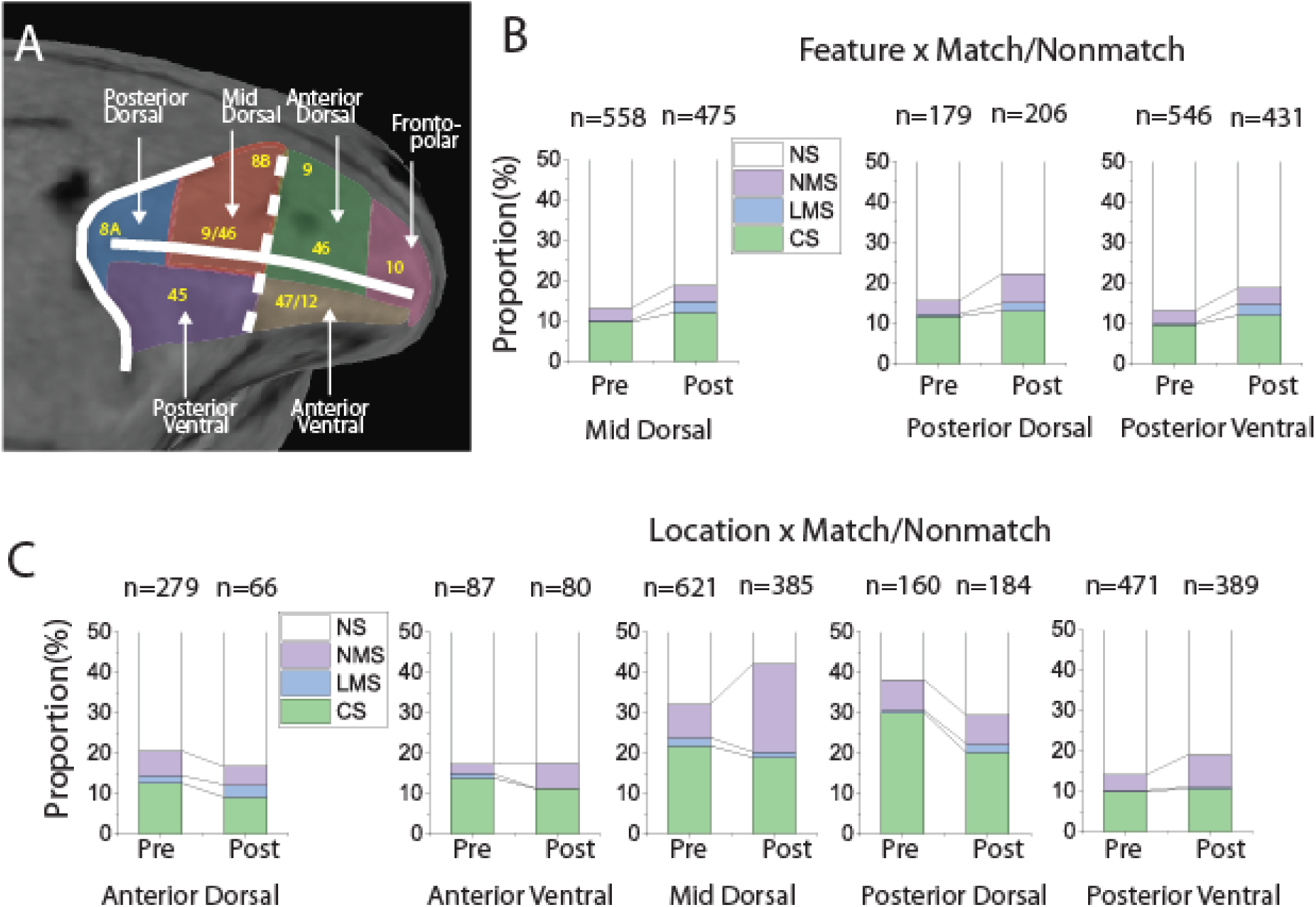
Cell selectivity changes by brain regions. (A) PFC subdivisions that were utilized for recording in the current study. (B) The effects of training in for the active feature task on the proportion of different selectivity categories (NS, CS, LMS and NMS) in the sample period. There were significant increases in the proportion of LMS cells in all three PFC regions included for analysis, but relatively low increases in the proportion of NMS cells. (C) The effects of training in the active spatial task on the proportion of different selectivity categories (NS, CS, LMS and NMS) in the sample period. The greatest increase in NMS occurred at the mid-dorsal region.

### NMS in task context

Previous theoretical studies linked NMS with more flexible readouts of multiple task variables, thus leading to the hypothesis that task complexity may modulate NMS. To test this hypothesis, we compared the neural responses to different shapes at the same location when the stimuli appeared as match or nonmatch in the conjunction task, to the same neurons’ responses to the same stimuli when they appeared in the feature task. In the conjunction task, animals needed to simultaneously remember both location and shape of visual stimuli, while in the feature task, they were only required to remember shape. Although the hypothesis predicted that the conjunction task would result in greater NMS than the feature task when the sensory stimuli was the same, this was not what we observed. No significant differences were observed for either CS cells (feature task sample proportion= 11.9%, conjunction task sample proportion=9.9%, exact matched pair sample proportion test, F=1.229, p=0.197, feature task delay2 proportion= 10.8%, conjunction task delay2 proportion=11.6%, exact matched pair delay2 proportion test, F=1.069, p=0.681) or NMS cells (feature task sample proportion= 4.1%, conjunction task sample proportion=4.4%, exact matched pair sample proportion test, F=1.031, p=0.901, feature task delay2 proportion= 4.6%, conjunction task delay2 proportion=5.9%, exact matched pair delay2 proportion test, F=1.278, p=0.266) (Fig. 6 A) in the sample period or the delay2 period that followed. We also examined changes in individual cells’ selectivity across the feature and conjunction tasks. Although this analysis was limited by the relatively low proportion of NMS cells in both tasks, we found an unstable mapping between tasks in the selectivity categories for both CS and NMS cells (Fig. 6 B), evidenced by the observation that most cells in CS and NMS category in the feature task changed their selectivity category in the conjunction task. There is a possibility, however, that the majority of the informative cells were simply due to chance (p=0.05 was used as threshold for detecting significant terms), which is enforced by the observation that NMS cells with larger degree of interaction in one task tend to also fall into the NMS category in the other task (Fig. 6 C). No significant difference was observed in the proportion of NMS cells when we performed a similar comparison between the spatial and conjunction task with a relatively small sample size (Fig. S1).

**Figure 6.**
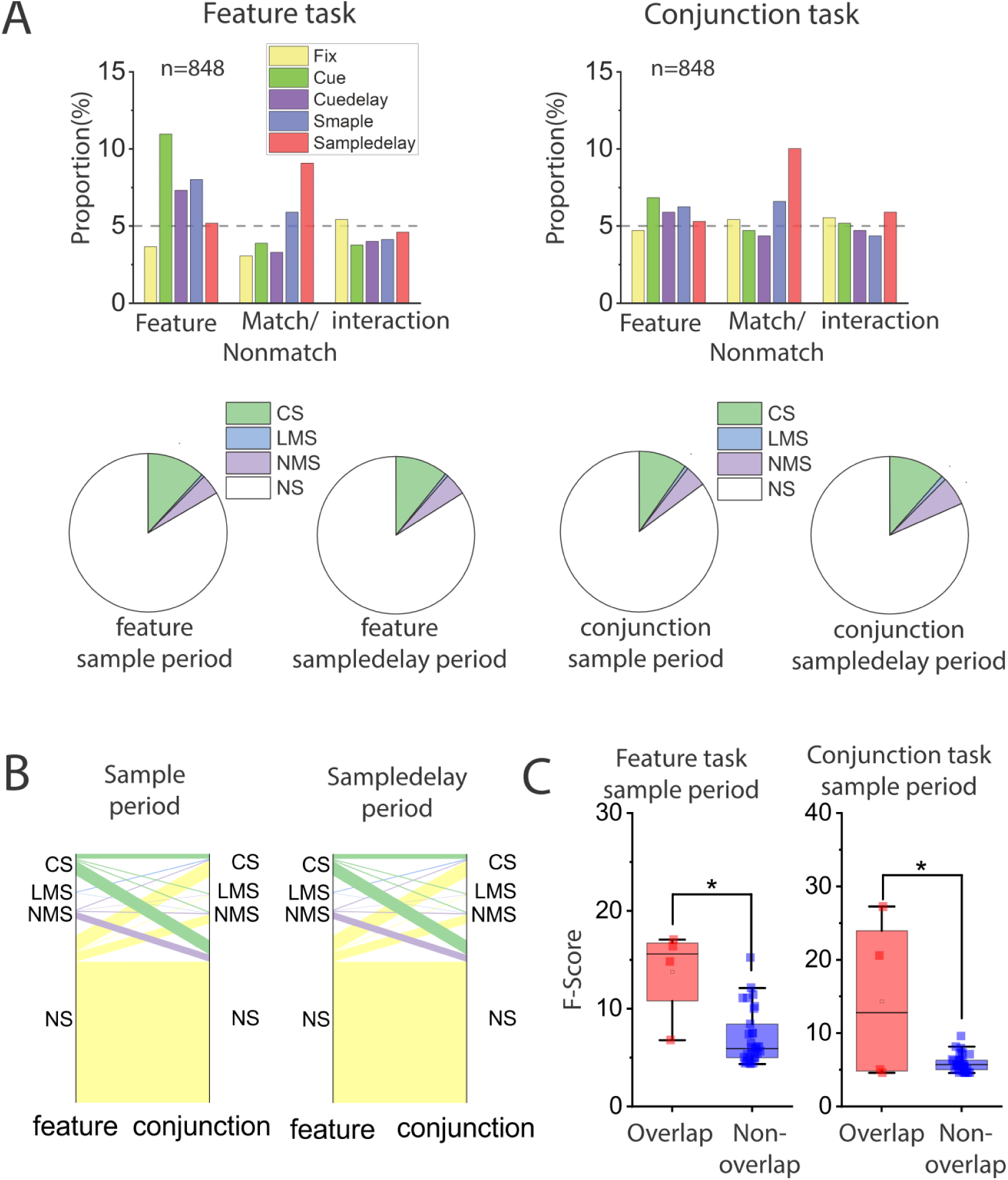
Cell selectivity in tasks with different task complexity. Analyses were performed on neural data from the same population of cells, with matching numbers of trials in the feature and the conjunction tasks. Only trials with the same stimuli were included in this analysis. (A) Examining interaction (NMS) across stimulus preference and matching status. Bar graphs show the proportions of cells tuned to stimuli shape, trials matching status and their interaction in different stages of both the feature and conjunction tasks. Pie charts display the proportion of different selectivity categories (NS, CS, LMS and NMS) in corresponding sample and delay2 periods. (B) Cell selectivity category mapping cross tasks. (C) F scores of the interaction term in the ANOVA were compared between cells that were classified as NMS cell in both tasks (overlapping cells), and those only classified as NMS in one of the tasks (non-overlapping cells).

The comparison of the naïve and trained conditions allowed us to test the overall incidence of NMS in different populations of PFC neurons, sampled randomly before and after training, which was carried out over the course of several months. If NMS were critical for the representation of task-relevant information, we would expect a difference in the observed proportion of neurons with NMS, when animals are passively viewing stimuli vs. when they are actively performing the task and storing representations of the stimuli in their WM. We therefore applied a two-way ANOVA to compare the neural responses of neurons between the active and passive spatial tasks after the monkeys had been trained to perform the active spatial task. We ultimately observed an increase in the proportion of cells that coded matching status during the sample period, as well as an increase in the proportion of cells coding sensory information in the delay1 period when the animal was prompted to report the matching decision. However, the observed increase in the proportion of NMS cells was not significant (passive proportion= 9.3%, active proportion=11.7%, exact matched pair proportion test, F=1.385, p=0.362) (Fig. 7 A). Interestingly, a large proportion of cells changed their selectivity category across tasks, especially for CS cells (Fig. 7 B), and the degree of NMS does not seem to be predictive of whether a given neuron would fall in the same selectivity category in both tasks (Fig. 7 C).

**Figure 7.**
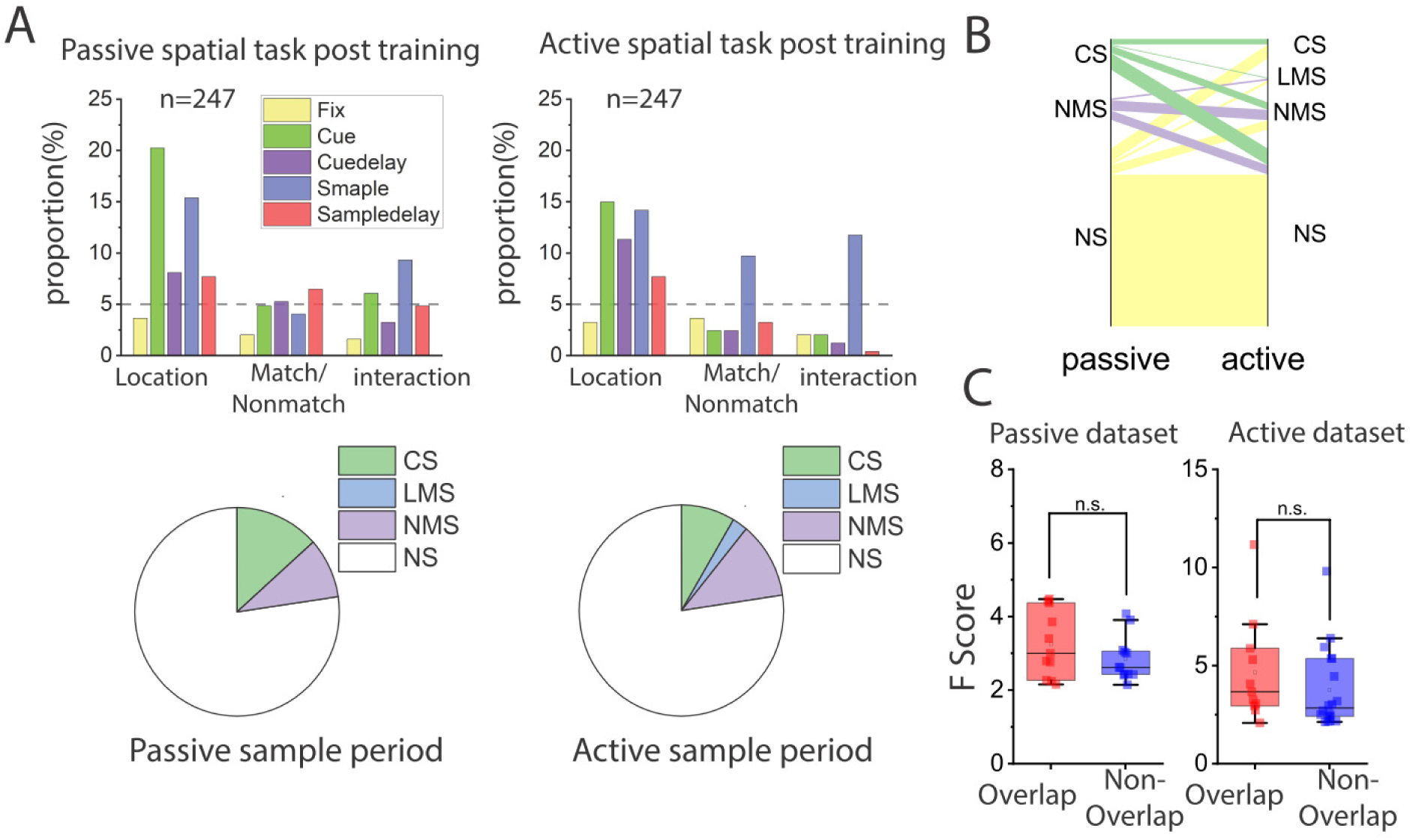
Cell selectivity in tasks with the same sensory input but different behavioral requirements. Analyses were performed on neural data from the same population of cells, with matching number of trials in the passive and active spatial tasks. Only trials that had the exact same stimuli pairs in both tasks were included in this analysis. (A) Examining interaction (NMS) between stimulus preference and matching status. Bar graphs show the proportions of cells tuned to stimuli location, trials matching status and their interaction in different stages of the tasks. Pie charts display the proportion of different selectivity categories (NS, CS, LMS and NMS) in the sample period. (B) Cell selectivity category mapping across tasks in the sample period. (C) F scores of the interaction term in the ANOVA were compared between cells that were classified as NMS cell in both tasks (overlapping cells), and those only classified as NMS in one of the tasks (non-overlapping cells).

### Information encoding by NMS neurons

It is known that training leads to increased incorporation of task relevant information in neural populations, with relatively little change to stimulus information (confirmed in our dataset, Fig. S2). However, the relative contribution by NMS is not clear. To quantify the amount of task relevant information contained in linear (CS and LMS) and NMS cells, we used a linear SVM decoder to decode sensory information (location and shape) and match/or nonmatch status information. Since the cell selectivity category could be defined by their response in either sample or delay2 period, we randomly selected equal numbers of linear and nonlinear cells in both task epochs for each comparison. The random selecting process was repeated multiple times to obtain a confidence interval. We ultimately found that linear and nonlinear cells contain comparable amounts of linearly decodable information in regard to both sensory information and task relevant information. The only observed difference between the decodable information in the linear and nonlinear cells occurred in the post training feature task, where linear cells were observed to contain more stimulus information in the sample period.

We also applied cross temporal decoding to compare classic and linear mixed cells in regard to population coding dynamics during the delay period. If information were represented by a stable pattern of activity, the classifier trained at one timepoint would be expected to work equally effectively at other time points where the information is present. Conversely, if information were represented by dynamic patterns of activity, then the decision boundary at one time point would not contribute to decoding information at other time points. The most prominent result from this analysis is that NMS cells produced significantly more stable code for matching spatial task information, compared to CS and LMS cells, as indicated by higher performance off the diagonal during the delay2 period (Fig. 9).

**Figure 8.**
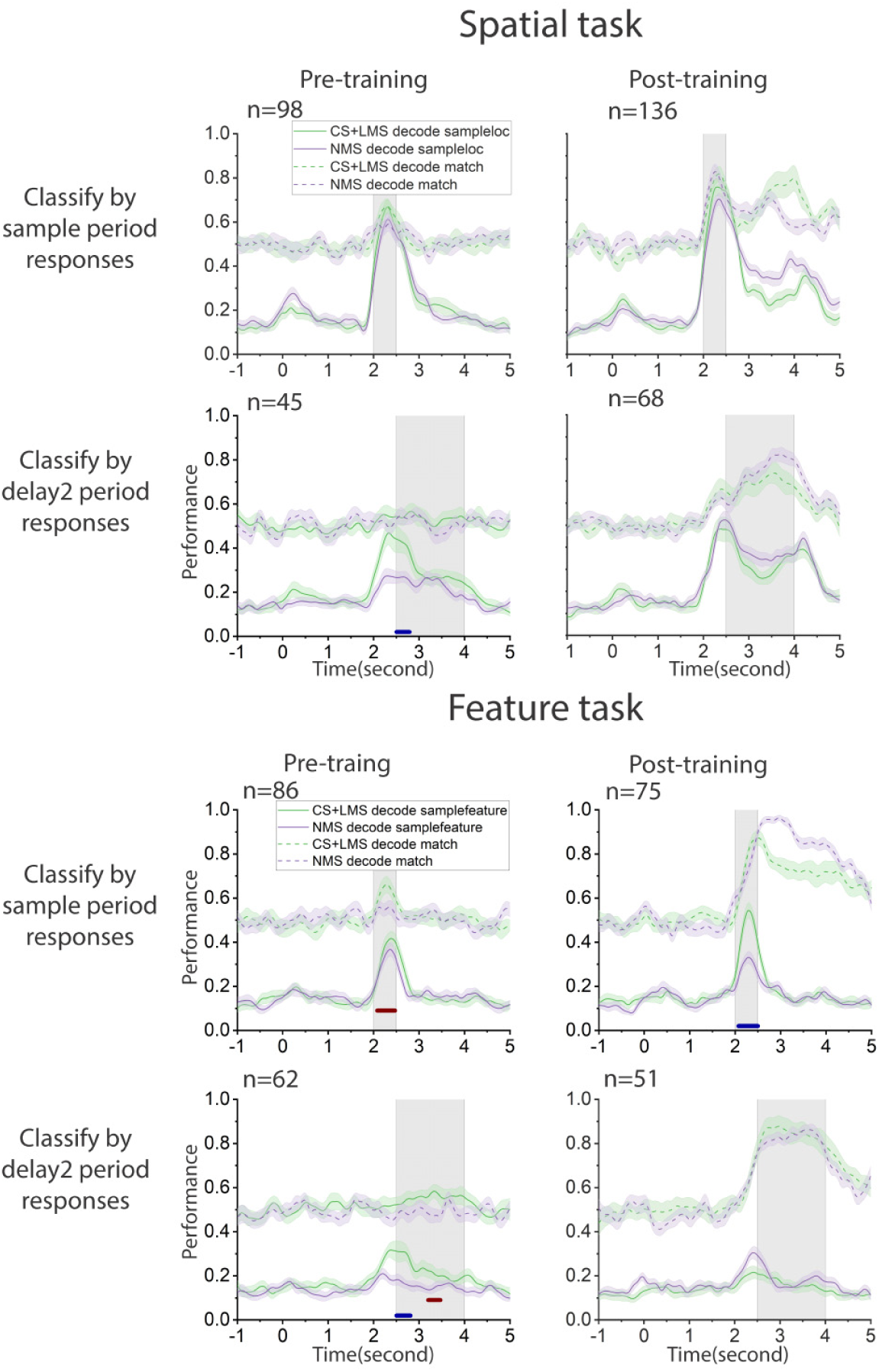
Comparing linear SVM decoder information between pure and linear selective cells (CS and LMS) vs. NMS cells. An equal number of linear (CS and LMS) and NMS cells were randomly selected from the sample or delay2 period. The selectivity categories were defined by spiking count in corresponding periods with reference to stimuli identity (stimuli location or shape) and matching status. The decoders were trained to classify either stimuli identity or match/nonmatch status with z-score normalized pseudo-population response. (A) Decoding performance in the spatial task before and after training. (B) Decoding performance in the feature task before and after training. Red and blue bars indicate time points when the performance for NMS and linear cells differs significantly within the shaded regions.

**Figure 9.**
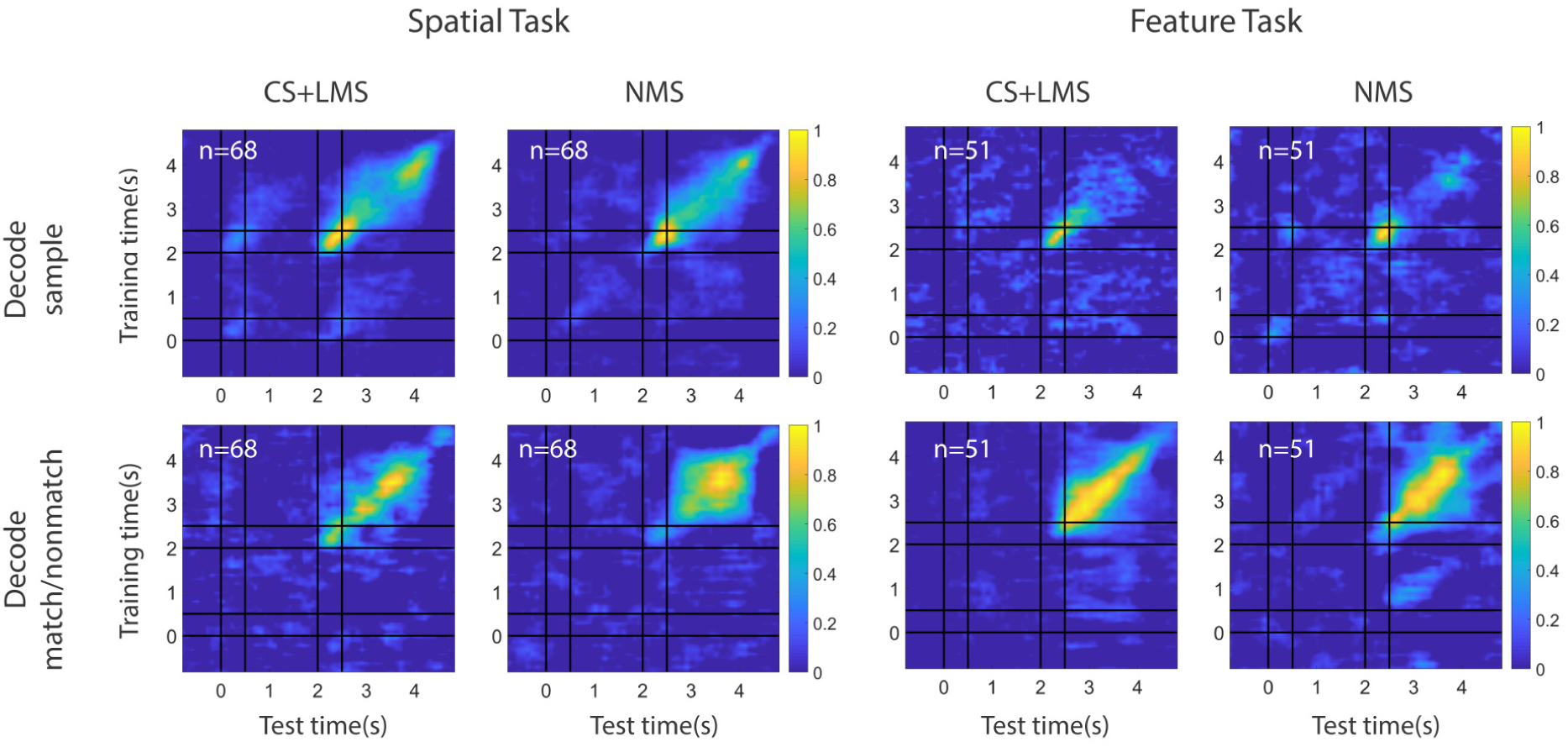
Coding dynamics of pure and linearly selective cells (CS and LMS) vs. NMS cells. Linear kernel SVM decoders were trained to perform cross-temporal decoding with different selectivity populations in the delay2 period, for both spatial and feature tasks, as indicated by the Y-axis. The decoder was then required to predict whether a match or non-match occurred at each time point based on a different test set of data, as indicated by the X-axis. Normalized decoding accuracy is indicated in the color bar, demonstrating how spatial and feature WM representations can be decoded from specific patterns of neural activity. Coding of matching information for NMS cells is more stable across time for the spatial task.

### NMS in correct and error trials

The presence of decodable information in the PFC does not necessarily imply the presence of information in the conscious mind, and the representation of task relevant information is ultimately revealed by the ability to conduct the task successfully. In order to decipher the role of NMS, we examined the F score of the main effects and their interaction in the ANOVA test in correct vs. error trials for the spatial task (Fig. 10), which displayed higher NMS levels than the feature or conjunction tasks. Similar to the pre- vs. post training comparisons, we examined two types of mixed selectivity: stimulus location vs. matching status and stimulus identity vs. task epoch. The number of trials and task variables were matched for each cell to avoid confounds in the comparison. The mean F score in correct trials for the location variable in the stimulus epochs for the location × epoch comparison was equal to 1.86, while for the error trials the mean was 2.59 (paired t test, t(147)=3.38, p=9.42 × 10^−4^). The effect extended into the delay epochs, where the average F score for location in correct trials was 1.43, and that of error trials was 1.79 (paired t test, t(150)=2.61, p=0.010). However, we did not find any differences in the F score for the interaction terms in any comparison. The results indicate that NMS may not be necessary in representing task relevant information in WM.

**Figure 10.**
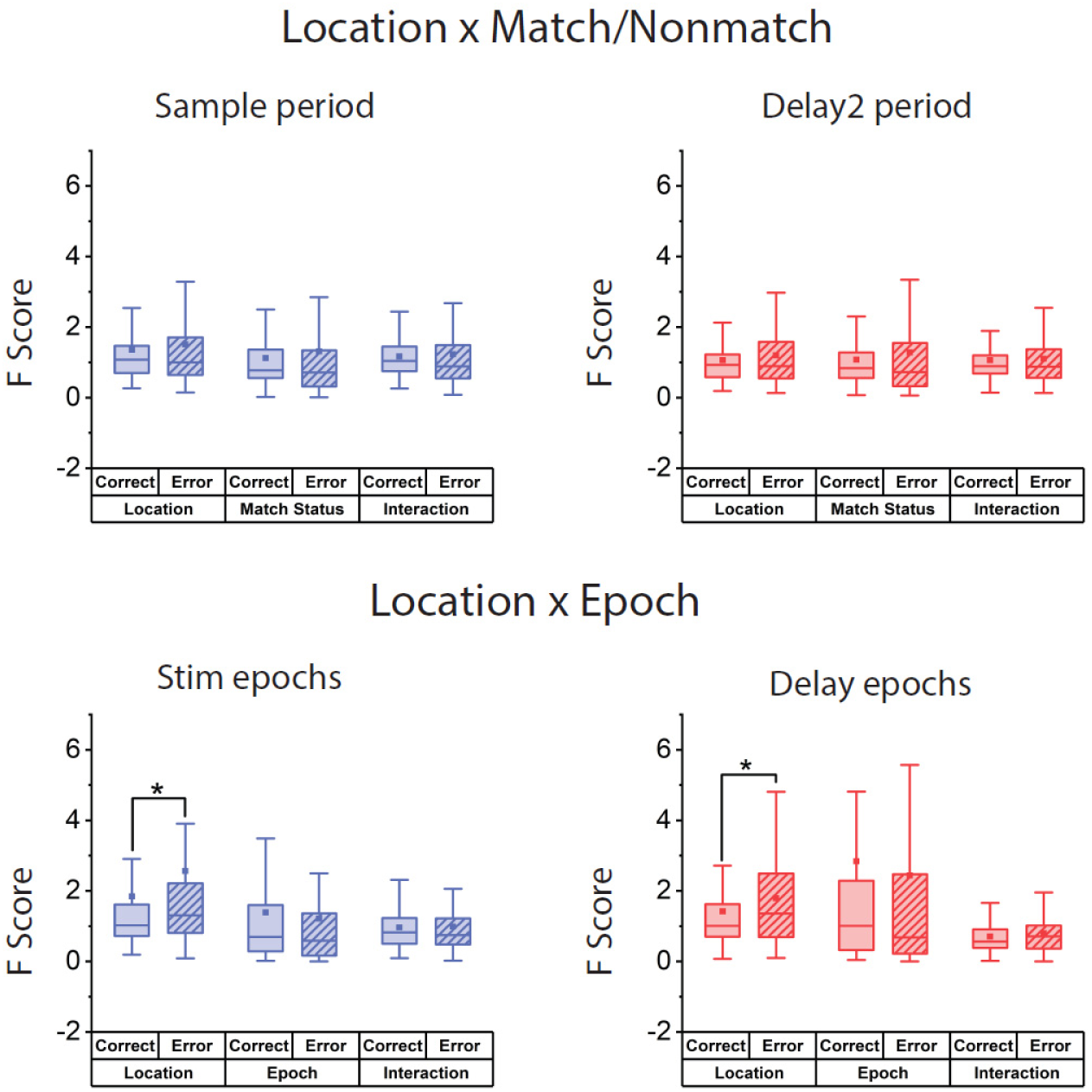
Comparison of cell selectivity in correct and error trials in the same population for the spatial task, after controlling for trial number and location pairs used. Two forms of mixed selectivity were examined (location x matching, location x task epoch). No change (location x matching delay2 period) or increase (location x task stim and delay epochs, location x matching sample period) in the F score of the interaction term of ANOVA results were observed in cells with significant interaction (NMS) terms. Higher F score for the variable of stimuli location was observed in error trials in the location x Epoch comparison. Box boundaries represent 25%-75% data range, whiskers indicate 1.5 IQR and squares indicates means across cells.

## DISCUSSION

Selectivity for different types of information is critical in representing the plethora of stimuli and task contexts that can be maintained in WM. NMS is thought to be critical in that respect, as it allows efficient representation of flexible, arbitrary combinations of variables (Barak et al., 2013; Buonomano and Maass, 2009; Fusi et al., 2016; Johnston et al., 2020; Rigotti et al., 2010). Consistent with this idea, increased dimensionality in NMS has been highlighted as a potential means of increasing the efficiency of WM task performance (Johnston et al., 2020; Rigotti et al., 2013) and dimensional collapse characterizes task errors (Rigotti et al., 2013). Moreover, all task relevant information could be decoded from NMS neurons alone, despite their relative scarcity, with decoder accuracy actually increasing as the task became more complex (Rigotti et al., 2013). NMS is assumed to emerge with training in complex tasks that combine multiple types of information, or in multiple tasks, even without an explicit requirement to combine such information (Johnston et al., 2020; Lindsay et al., 2017). However, this idea has not been tested experimentally until now. Our study, by virtue of analyzing neural recordings before and after training in a series of cognitive tasks, directly tested these postulates. We found that NMS resulted in a modest increase with training, but only for some tasks, and furthermore, task complexity was not a predictor of NMS emergence. A causal relationship between success and dimensionality—and by extension, NMS—was also not supported by our results, as we did not observe any significant changes in NMS between error and success trials. These insights refine and qualify the role NMS plays in WM, and identify a number of open questions.

### Effects of training on neural responses

WM is considerably plastic and at least some aspects of it, such as mental processing speed and the ability to multitask, can be improved with training (Bherer et al., 2008; Dux et al., 2009; Jaeggi et al., 2008; Klingberg et al., 2005; Klingberg et al., 2002). WM training has been proven particularly beneficial for clinical populations, e.g. in the case of traumatic brain injury, attention deficit hyperactivity disorder (ADHD), and schizophrenia (Klingberg et al., 2002; Subramaniam et al., 2012; Westerberg et al., 2007). However, the verdict of whether WM training confers tangible benefits on normal adults and whether these benefits transfer to untrained domains, remains a matter of heated debate. (Constantinidis and Klingberg, 2016; Cortese et al., 2015; Fukuda et al., 2010; Owen et al., 2010; Peijnenborgh et al., 2015; Schwaighofer et al., 2015).

This malleability of cognitive performance is thought to be mediated by the underlying plasticity in neural responses, most importantly within the PFC (Constantinidis and Klingberg, 2016). In a series of prior studies, we have investigated changes in PFC responsiveness and selectivity (Meyer et al., 2011; Meyers et al., 2012; Qi et al., 2011; Riley et al., 2018), as well as other aspects of neuronal discharges such as trial-to-trial variability and correlation between neurons (Qi and Constantinidis, 2012a, 2012b). This led to our present analysis where—guided by experimental and theoretical predictions (Rigotti et al., 2013)—we examined NMS as another potential source of enhanced ability to represent WM information after training.

In agreement with our hypothesis, we found that training increased the proportion of neurons that exhibit NMS. However, training does not seem to be a prerequisite, as NMS was also observed in animals that were naïve to any cognitive training. Prior research has established that the human and primate PFC represent stimuli in memory even when not prompted to do so (Foster et al., 2017), or without training in WM tasks (Meyer et al., 2007). Our finding of NMS neurons in naïve monkeys provides another exemplar of that principle. However, NMS only increased for certain types of task information and not for others, thus suggesting that its role in the simultaneous representation of multiple types of information is not universal across tasks. These results stand in agreement with some prior studies, which have failed to uncover substantial NMS in the tasks they employed (Cavanagh et al., 2018). Examining where NMS failed to appear—and where WM representations fail to spontaneously appear— will be an important area of future investigation for NMS.

The greatest increase in the proportion of neurons that exhibited NMS for the spatial working memory task was observed at the mid-dorsal region during the sample presentation period (Fig. 5). This disproportionate increase in NMS neurons was associated with a modest decrease in neurons that exhibit CS, as predicted by theoretical studies (Lindsay et al., 2017). However, this finding, too did not generalize across conditions. During the delay periods of the spatial task, we saw an across-the-board increase in neurons with CS, which were much more abundant in the trained than the naïve PFC (Fig. 3). In fact, the increase in feature selectivity after training was driven almost exclusively by CS cells, suggesting a potential division of labor between NMS and CS in the PFC for different types of information.

### Task Complexity and Difficulty

Another potential factor that determines the emergence of NMS is task complexity. NMS may arise exclusively in highly complex tasks that require subjects to maintain and combine multiple types of information in their WM, simplifying the involved neural circuits to achieve greater efficiency (Rigotti et al., 2013). We thus tested this concept by applying a dataset that relied on three tasks which differed in complexity (and overall difficulty). The spatial and feature tasks each required maintenance of a single stimulus property in memory (location or shape). The conjunction task required both. Surprisingly however, we did not observe a higher incidence of NMS in the conjunction task when compared to the feature task. Moreover, we observed a much lower incidence of NMS in the feature task compared to the spatial task despite the fact that the latter was no more complex or difficult for the monkeys to perform (Meyer et al., 2011; Riley et al., 2018). This implies that NMS in the PFC may not be necessary for certain types of information, like object shape, even when the task complexity is high. Alternatively, we may also consider the possibility that the increased NMS that results from training may be sufficient for the majority of behavioral requirements, without any additional increases required. Future research is therefore necessary to assess and examine all of these possibilities, and more.

### Regional Specialization

Different types of information are represented across the dorso-ventral and anterior-posterior axes of the PFC (Constantinidis and Qi, 2018), and examining the regional distribution of NMS neurons within the PFC therefore bears a clear importance. We found that NMS was most strongly demonstrated in the mid-dorsal area for the spatial task and the posterior dorsal area for the feature task. This pattern was generally consistent with the known distribution of neuronal selectivity for stimuli in the PFC (Riley et al., 2018).

### Information Content and Task Performance

A critical issue regarding the role of Mixed Selectivity is whether nonlinear match selectivity, by virtue of representing information more efficiently, is also more necessary for effective task performance (Rigotti et al., 2013). We relied on a linear SVM decoder to decipher the specific information that may be represented by NMS cells, compared to CS cells. In the current study, we found similar quantities of information could be decoded from (equal-sized) populations of CS and NMS neurons, though the coding dynamics for some types of information were significantly different between CS and NMS cells. Similarly, when we compared the NMS levels of successful and failed task trials, we were surprised to find that there was no appreciable difference. This suggests that loss of information encoded in a nonlinear manner is not the primary factor of successfully maintaining information in the conscious mind. An important caveat for this conclusion is that the combination of small and unbalanced number of error trials in match vs. nonmatch conditions make the detection power for the interaction fairly small in our analysis. Moreover, with very little NMS presented even in success trials after matching the trial number for the error condition, a floor effect may have prevented a further decline from becoming apparent. Nonetheless, our result reinforces the idea that NMS is not necessary in all tasks, without which performance fails. An interesting observation in this analysis was that the proportion of cells tuned to spatial location was elevated in error trials. The result may imply that task success also depends on the task relevance of the represented information, with error trials incorporating greater quantities of task irrelevant spatial information and therefore unnecessarily drawing away WM resources without benefit. Ultimately, by comparing and evaluating the conditions in which NMS emerges, we may decipher its true role in WM and other cognitive functions.

## ACKNOWLEDGMENTS

Research reported in this paper was supported by the National Eye Institute of the National Institutes of Health under award number R01 EY017077. We wish to acknowledge Kathini Palaninathan and Austin Lodish for technical help; Junda Zhu, and Sihai Li for helpful comments on the manuscript.

The authors report no conflicts of interest during the research conducted.

---

**Fig. S1.**
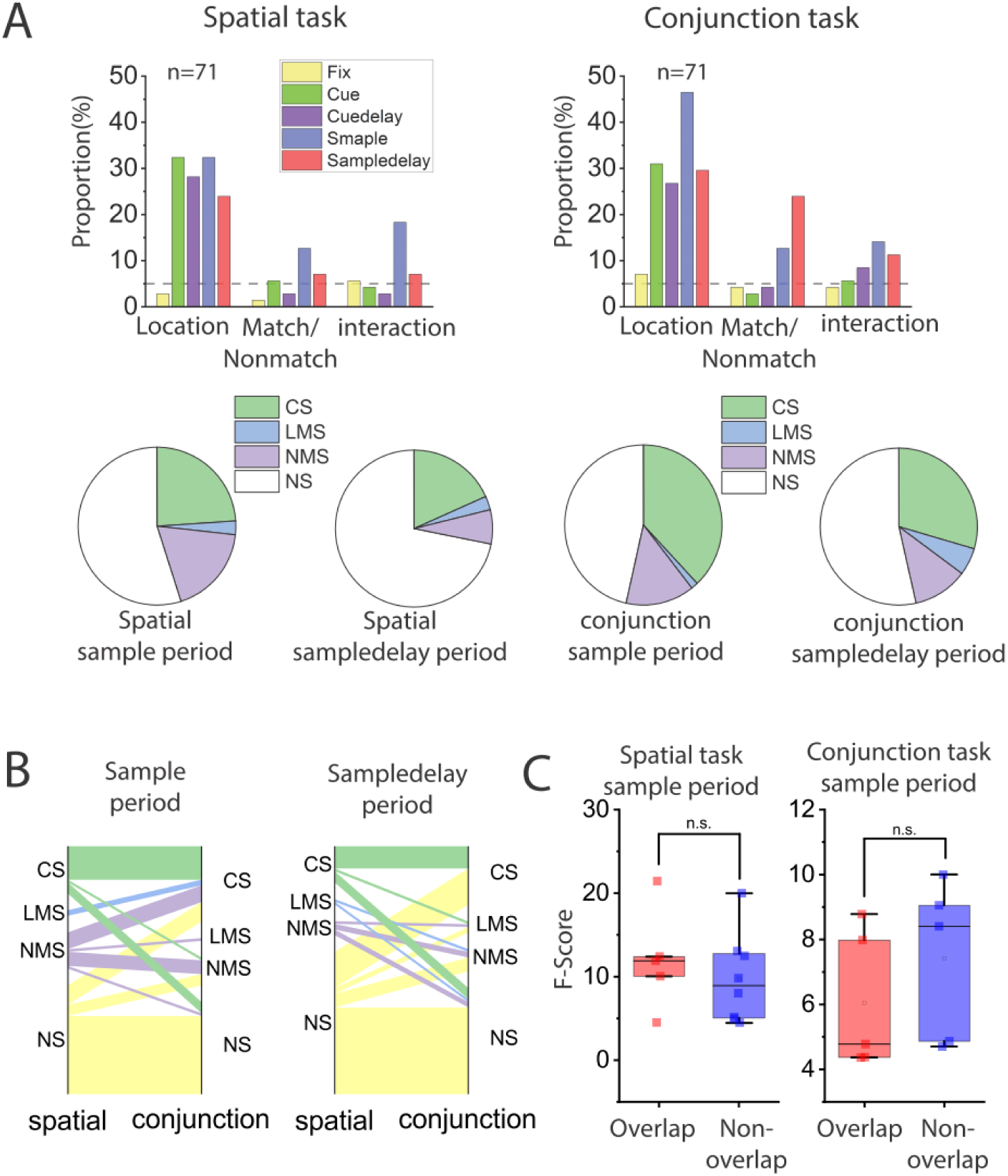
Plotting conventions same as Fig. 6. Comparing cell selectivity between the spatial and the conjunction tasks. The analysis was performed on neural data from the same population of cells, with matching numbers of trials in the spatial and the conjunction tasks. Only trials with the same stimuli were included in this analysis. (A) Examining interaction (NMS) between stimulus preference and matching status. Bar graphs display the proportions of cells tuned to stimuli shape, trials matching status and their interaction in different stages of the tasks, in both the feature and conjunction task. Pie charts display the proportion of different selectivity categories (NS, CS, LMS and NMS) in corresponding sample and delay2 periods. (B) Cell selectivity category mapping across tasks. (C) F scores of the interaction term in the ANOVA were compared between cells that were classified as NMS cells in both tasks (overlapping cells), and those only classified as NMS in one of the tasks (non-overlapping cells). Similar to the comparison between the feature and the conjunction tasks, no change in the proportion of NMS was observed.

**Fig. S2.**
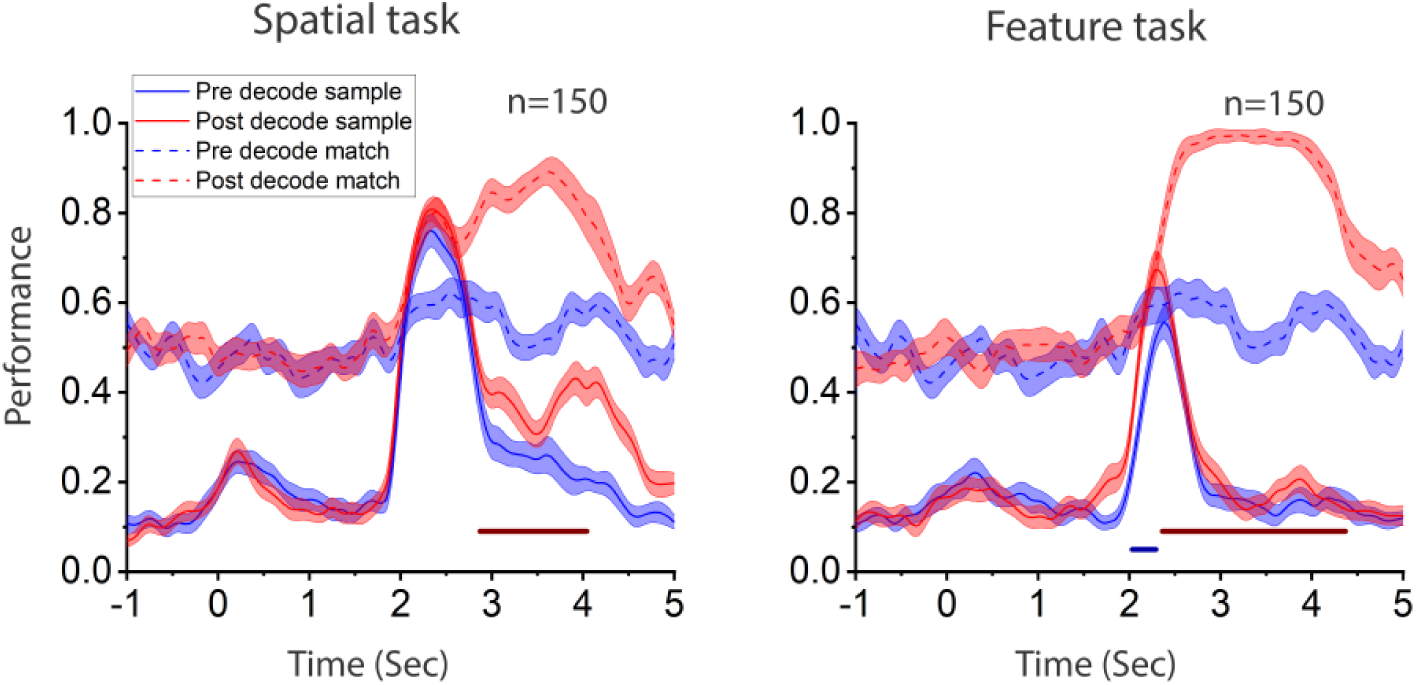
Incorporation of new information after training for the spatial and the feature tasks. Linear SVM decoders were trained to classify either stimuli identity or match/nonmatch status with z-score normalized pseudo-population response. Color bars indicate time points when the performance for NMS and linear cells differs significantly. Red bar for decoding matching status, blue bar for decoding stimuli identity.

